# The crystal structure of the naturally split gp41-1 intein guides the engineering of orthogonal split inteins from a *cis*-splicing intein

**DOI:** 10.1101/546465

**Authors:** Hannes M. Beyer, Kornelia M. Mikula, Mi Li, Alexander Wlodawer, Hideo Iwaï

## Abstract

Protein *trans*-splicing catalyzed by split inteins has increasingly become useful as a protein engineering tool. The 1.0 Å-resolution crystal structure of a variant from naturally split gp41-1 intein, identified from the environmental metagenomic sequence data, revealed an improved pseudo-C2-symmetry commonly found in the Hedgehog/Intein (HINT) superfamily with extensive charge-charge interactions between the split N-and C-terminal intein fragments. We successfully created orthogonal split inteins by engineering a similar charge network in the same region of a *cis*-splicing intein. The same strategy could be applicable for creating novel natural-like split inteins from other, more prevalent *cis*-splicing inteins.

## Introduction

Protein splicing is a posttranslational modification where an intervening protein (intein) residing within an unrelated host protein excises itself, thereby covalently ligating the N-and C-terminally flanking sequences (exteins) with a standard peptide bond^1,2,3,4^. As a result, the ligated product is scar-less and devoid of any indication of its previous merged existence, while the function of the host protein is generally restored (Fig. 1). Inteins are commonly regarded as selfish parasitic elements, albeit some evidence attributes a regulatory role in controlling the activity of host proteins in response to environmental cues triggering the splicing reaction^3,5^. During recent years, inteins have become increasingly popular for diverse applications in biotechnology, chemical biology, and synthetic biology because of the following properties. First, intein-mediated protein splicing tolerates the deliberate exchange of extein sequences^3,6^. Second, the existence of naturally occurring split inteins reconstituting a functional protein from two polypeptide chains, as well as the possibility of splitting *cis*-splicing inteins, generates ample possibilities for applications with protein *trans*-splicing (PTS)^3,7,8^ (Fig. 1B). Ever since the discovery of protein splicing by inteins, engineering of inteins toward high performance, high tolerance of junction sequences, and smaller variants has been an ongoing quest^8,9^. Successfully engineered inteins arose from the accumulation of beneficial mutations upon directed evolution^10,11,12,13^, propagation of consensus sequence^14^, and as a result of rational design^15,16,17^.

**Fig. 1.**
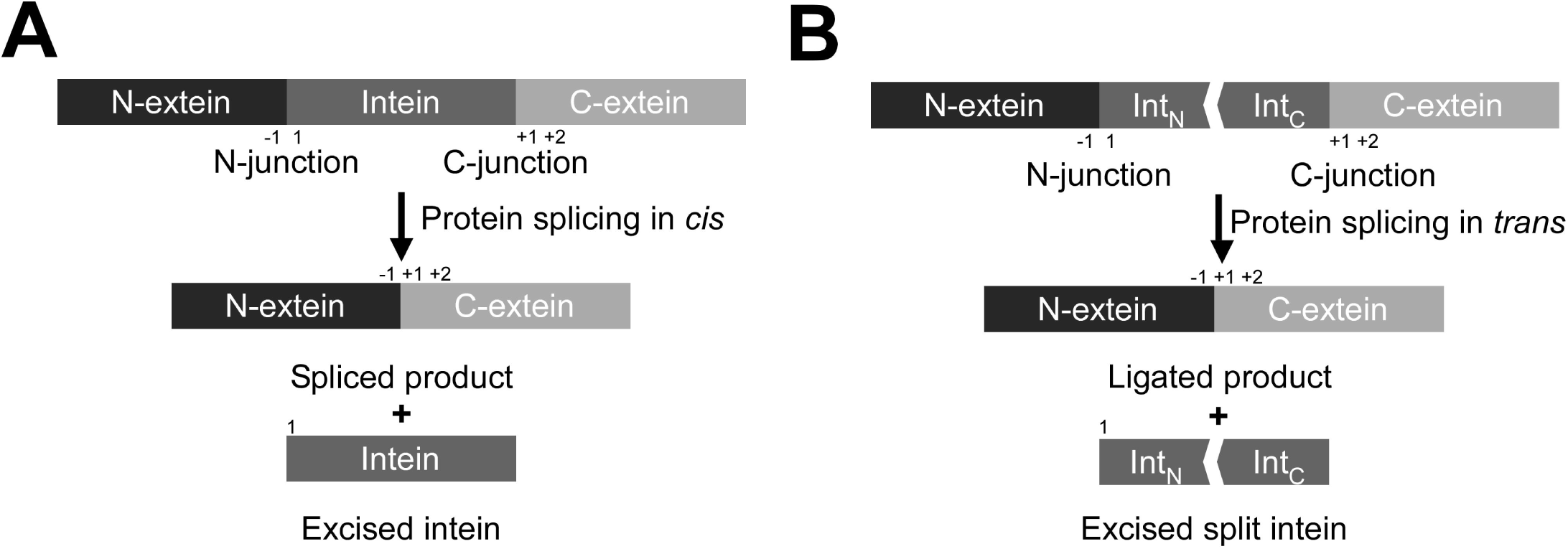
Schematic representation of protein splicing in *cis* and *trans*. (**A**) C*is*-splicing inteins excise themselves from a precursor where the N-and C-exteins flank the intein on the same polypeptide. (**B**) Protein *trans*-splicing (PTS) ligates N-and C-exteins, each originating from an independent polypeptide, with a covalent peptide bond resulting in a ligated product. Interaction of the N-(Int_N_) and C-terminal (Int_C_) split intein halves initiates the protein-splicing reaction. The N-terminal junction sequence at the front of an intein is termed as the -1 position. The +1 position after the intein sequence usually has Cys, Ser, or Thr residue. The second residue after an intein is numbered as the +2 position.

The naturally fragmented gp41-1 intein was found from metagenomic sequencing^18^. It is one of the smallest reported inteins with very fast splicing activity^19^, consisting of 88-residue N-terminal (Int_N_) and 37-residue C-terminal (Int_C_) fragments. Its small size and robust protein splicing activity make it an attractive template for protein engineering^19^. Also, gp41-1 intein has Ser as a catalytic residue at the +1 position. Given the much higher frequency of Ser over Cys within pro-and eukaryotic proteins, inteins with +1Ser allow a broader spectrum of possible insertion sites for scar-less protein ligation than naturally split inteins with Cys at the +1 position, thereby expanding potential applications.

Despite increasing interests in the utilization of various split inteins for protein engineering purposes, the repertoire of split inteins with both robust protein splicing activity and high sequence tolerance at the splice junctions are still small. Particularly, pairs of orthogonal split inteins are desirable for one-pot multiple fragment protein ligation by PTS requiring two orthogonal split inteins^20,21,22,23^. Previous engineering attempts to derive novel split inteins from naturally occurring *cis*-splicing inteins did not result in highly robust split inteins, indicating that *cis*-splicing inteins are not optimized for *trans*-splicing, unlike naturally occurring split inteins^15,24,25^.

Here we report the 1.0 Å-resolution crystal structure of the naturally split gp41-1 intein. Based on the crystal structure, we grafted the features found in the gp41-1 intein onto *cis*-splicing inteins to develop novel split inteins and demonstrated the engineering of orthogonal split intein fragments from a *cis*-splicing intein.

## Results

### Crystal structure of the gp41-1 intein

As the first step to engineer inteins based on the gp41-1 intein, we created a *cis*-splicing gp41-1 intein variant by genetically fusing the gp41-1_N_ and gp41-1_C_ split fragments. We found that the *cis*-splicing gp41-1 intein retained high protein splicing activity when the native three extein residues were kept (Fig. 2). Next, for structure determination, we crystallized an inactive mutant of the *cis*-splicing gp41-1 intein bearing an alanine mutation at the first residue (C1A). We solved the crystal structure of that inactive variant of the gp41-1 intein at the resolution of 1.0 Å by molecular replacement using the crystal structure of *Npu*DnaE intein as a search model (Table 1). The structure of gp41-1 has the canonical intein horseshoe shape, termed HINT (Hedgehog/INTein) fold (Fig. 2A). A Dali server search identified the engineered *Npu*DnaB^Δ290^ mini-intein (PDB ID: 4or1) as the closest structure to the gp41-1 intein with a Z-score of 20.1 and an r.m.s.d. of 1.4 Å for 127 residues^26^. *Npu*DnaB^Δ290^ intein is composed of 139 residues, which is 14 residues larger than the gp41-1 intein^15^. The main differences in the length between the two structures can be attributed to two distinct regions (Fig. 2B and 2C). One is in the split fragment-connecting loop where canonical inteins harbor a homing endonuclease domain insertion (C36 site)^25^. The other is a loop at the pseudo-C2-symmetry related site (N35 site)^25^. These regions account for 11 residues of the size difference.

**Table 1.** Data collection and refinement statistics.

|  |  |
| --- | --- |
| **Data collection** | Diamond i02 |
| Wavelength | 0.9795 |
| Space group | *I*222 |
| Molecules/a.u. | 1 |
| Unit cell *a*, *b*, *c* (Å);<br>$\alpha=\beta=\gamma$ (°) | 48.81, 69.99, 71.24<br>90, 90, 90 |
| Resolution (Å)* | 49.92-1.02 (1.04-1.02) |
| $R_{\text{merge}}$ (%)<sup>†</sup> | 3.8 (28.8) |
| $R_{\text{pim}}$ (%)<sup>‡</sup> | 1.9 (23.1) |
| No. of reflections<br>(measured/unique) | 242564/55708 |
| $\langle I/\sigma I \rangle$ | 28.3 (2.8) |
| Completeness (%) | 89.4 (37) |
| Redundancy | 4.35 |
| **Refinement** |  |
| Resolution (Å) | 49.92-1.02 |
| No. of reflections<br>(refinement/ $R_{\text{free}}$) | 53080/2642 |
| $R / R_{\text{free}}$ <sup>‡</sup> | 12.10/15.00 |
| No. atoms |  |
| Protein | 1093 |
| Ligand/ion | 51 |
| Water | 220 |
| R.m.s. deviations from ideal |  |
| Bond lengths (Å) | 0.020 |
| Bond angles (°) | 1.98 |
| Ramachandran plot |  |
| Favored (%) | 98.4 |
| Allowed (%) | 1.6 |
| PDB code | 6qaz |

\*The highest resolution shell is shown in parentheses.
<sup>†</sup> $R_{\text{merge}} = \sum_h \sum_i |I_i - \langle I \rangle| / \sum_h \sum_i I_i$ , where $I_i$ is the observed intensity of the $i$ -th measurement of reflection $h$ , and $\langle I \rangle$ is the average intensity of that reflection obtained from multiple observations.
<sup>‡</sup>Defined in Weiss et al., 2001<sup>52</sup>.
<sup>‡</sup> $R = \sum ||F_o| - |F_c|| / \sum |F_o|$ , where $F_o$ and $F_c$ are the observed and calculated structure factors, respectively, calculated for all data. $R_{\text{free}}$ was defined in Brünger, 1992<sup>53</sup>.

**Fig. 2.**
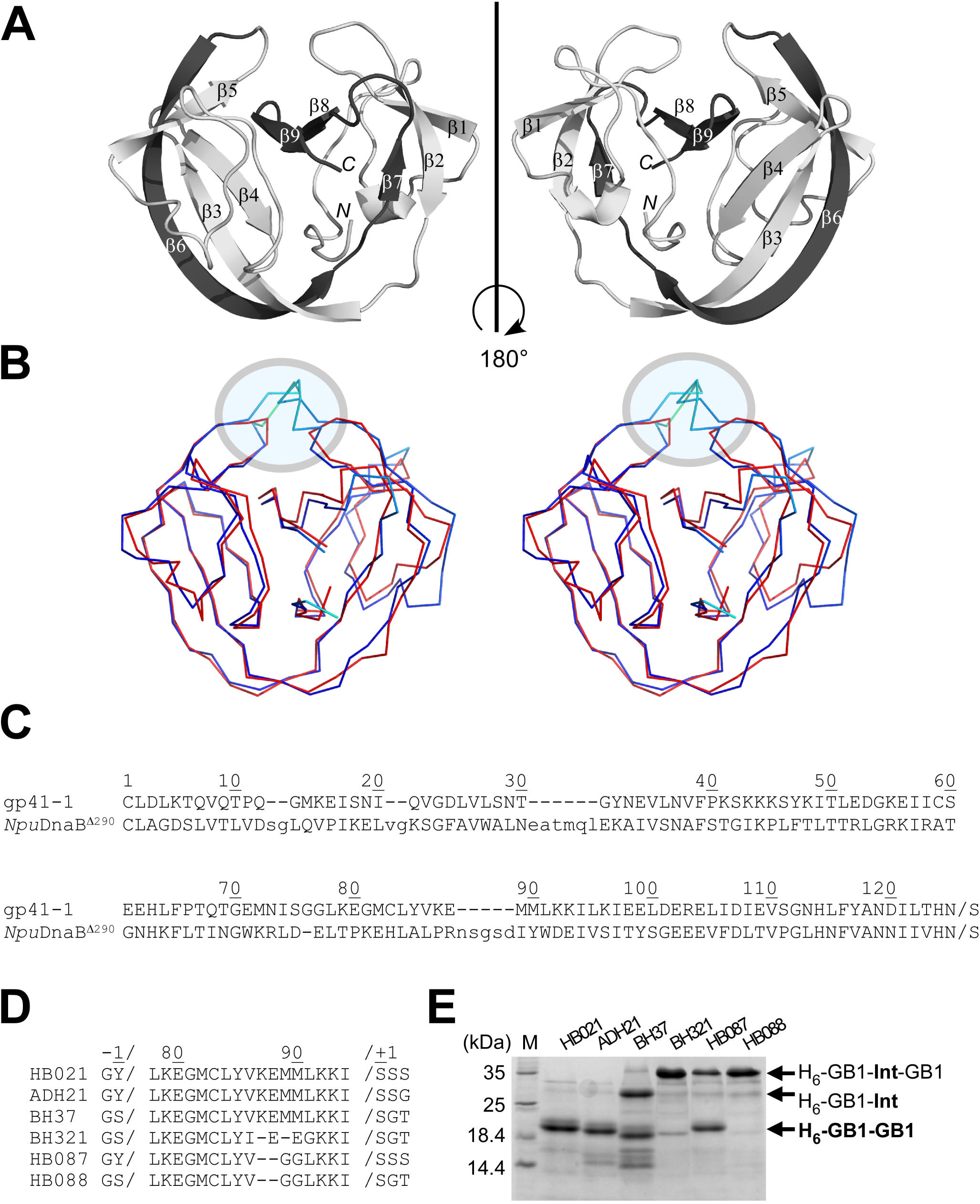
Crystal structure of the engineered *cis*-splicing gp41-1 intein. (**A**) Ribbon representations of the structure of gp41-1 intein. The region corresponding to the C-terminal split fragment (Int_C_) is colored in dark grey. *N* and *C* indicate N-and C-termini, respectively. (**B**) Stereo-view of an overlay of the crystal structures of gp41-1 intein (red) and the closest structure of *Npu*DnaB mini-intein (blue) (PDB: 4o1r). The major insertion sites in the *Npu*DnaB mini-intein are indicated by a circle. (**C**) Sequence alignment between the *cis*-splicing gp41-1 intein and *Npu*DnaB mini-intein (*Npu*DnaB^Δ290^). (**D**) Engineering of the gp41-1 intein in the loop and splicing junction regions and their effects on the protein splicing in *cis*. HB021, ADH21, BH37, BH321, HB087, and HB088 indicate the short names for different constructs with the sequence variations shown in the sequence alignment. (**E**) SDS-PAGE analysis of the *cis*-splicing activity of the engineered gp41-1 intein variants. M stands for molecular weight markers. H_6_-GB1-Int-GB1 indicates the unspliced precursor protein bearing variants of the intein. H_6_-GB1-GB1 indicates *cis*-spliced products with various junction sequences causing minor variations in the migration profile. H_6_-GB1-Int indicates an off-pathway cleavage product. Full-length gels are presented in Supplementary Figure 1.

The smaller size of the gp41-1 intein improved the symmetry of the pseudo-C2-symmetric structure found in the HINT fold by shortened insertions^27^ (Fig. 2). The two C2 symmetry-related regions (residues 3-52 and residues 60-110) can be well superimposed^27,25^ (Fig. 3A). The gp41-1 intein structure can be thus dissected into four distinct units: the first C2-symmetry related unit, β-strand (β4), the second C2-symmetry related unit, and two β-strands (β8, β9) (Fig. 3B). The C2-symmetry related unit can be further divided into a globular region and two β-strands (β2 and β3, or β6 and β7). The naturally split site of the gp41-1 intein is located within the second C2-symmetry related unit, separating the C2-symmetry unit into a globular region and two β-strands (Fig. 3D).

**Fig. 3.**
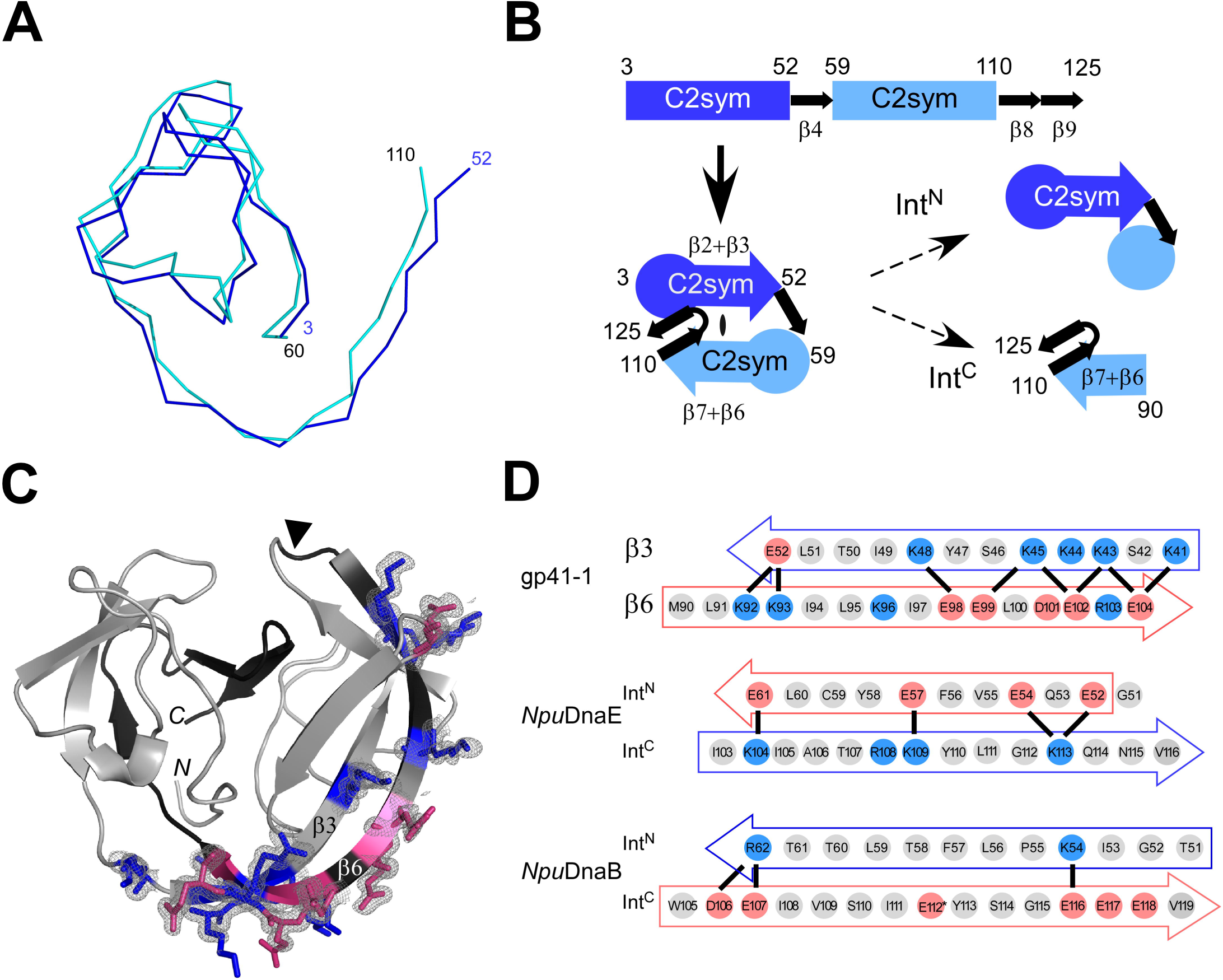
The modular architecture of the gp41-1 intein and the charge network. (**A**) An overlay of the backbone atoms of the two C2-symmetry related units (residues 3-52 and 59-110) observed in the gp41-1 structure. (**B**) Arrangement of the C2-symmetry related units and connections of the secondary structures. The natural split site of the gp41-1 intein locates within the second C2-symmetry related part and at the front of β6 strands. (**C**) The charged network found in the gp41-1 intein structure. The side-chains of the charged residues in β3 and β6 strands are shown together with the electron densities. Residues with negative and positive charges are highlighted in red and blue, respectively. The natural split site is indicated by a filled triangle. N and C indicated the N-and C-termini, respectively. (**D**) Comparison of the charged residues in β3 and β6 strands between the gp41-1, *Npu*DnaE, and *Npu*DnaB inteins. Thick lines indicate possible favored charge interactions. An asterisk indicates Glu112 modeled as Val112 in the coordinate of the *Npu*DnaB mini-intein structure (PDB: 4o1r).

### Minimizing gp41-1 intein

The first question we asked was whether it is possible to minimize the *cis*-splicing gp41-1 intein to a size even smaller than 125 residues. The conserved insertion site for a homing endonuclease found in canonical inteins also overlaps the split sites most commonly found for many split inteins^25^. We removed two residues from the linker where Int_C_ and Int_N_ were connected, i.e., at the natural split site of the gp41-1 intein. This deletion drastically reduced the protein-splicing activity (Fig. 2D). Our attempt at optimizing the linker sequence to rescue the robust splicing activity of the gp41-1 intein was unsuccessful (Fig. 2D). Whereas the closest crystal structure of *Npu*DnaB^Δ290^ intein shows higher B-factors for the backbone atoms of the corresponding linker region (43.4 ± 2.7 Å^2^), the B-factors for the corresponding regions in the gp41-1 intein crystal structure are much lower (22.8 ± 3.1 Å^2^), suggesting less flexibility and more ordered structure in this region of the gp41-1 intein. This region also contains an unusual *cis*-peptide bond between Lys87 and Glu88 – its presence is unambiguously supported by the excellent electron density, although part of the side chain of Glu88 appears to be disordered. The gp41-1 intein does not seem to tolerate any deletion easily. The linker length at this site could play an essential role in the productive folding of some HINT superfamily members^28^. Moreover, we observed a drastic reduction of protein splicing activity when deviating the extein sequences from the native sequence (Y-1, S+2, and S+3) (Fig. 2D). Thus, gp41-1 intein might not be a suitable intein when the natural junction sequences require modifications.

### The charge network in the gp41-1 intein

Previously, it has been suggested that local charge distributions between naturally split intein halves are important for their association^18^, ^23,29^. We observed extensive charge-charge interactions in the crystal structure of the gp41-1 intein, as observed among other naturally split inteins^29^. Particularly, they are located in the interacting regions within the β-strands of the two C2-symmetry related units. Recently, a “capture and collapse” model has been proposed as a folding mechanism for the naturally split DnaE intein from *Nostoc punctifome* (*Npu*DnaE intein), in which the first step of the interaction between the split fragments is initiated by electrostatic interactions on the extended β-strands^30^. We identified similar electrostatic networks between β6 at the beginning of Int_C_ and β3 at the C-end region of Int_N_ in the structure of the gp41-1 intein (Fig. 3C and 3D). These two anti-parallel β-strands appeared to form a charge zipper, reminiscent of leucine zipper structures but embedded in extended strands rather than helices^31^. We also compared the charge patterns in the same regions with the naturally split *Npu*DnaE intein as well as with the closest structural homolog, the *cis*-splicing *Npu*DnaB mini-intein (Fig. 3D). Whereas the *Npu*DnaB mini-intein does not contain such an extensive charge network in the corresponding region, the gp41-1 intein encompasses more prominent charge interactions than the *Npu*DnaE intein (Fig. 3D). This observation might support the notion that the “capture and collapse” model suggested for the naturally split *Npu*DnaE intein might also be valid for the naturally split gp41-1 intein^30^.

Interestingly, the Int_C_ region of the gp41-1 intein is dominated by negative charges, as opposed to the more positive charges found in the same region of *Npu*DnaE intein (Fig. 3D). The charge distributions in the β3 and β6 are thus opposite between the naturally split *Npu*DnaE and gp41-1 inteins (Fig. 3D). Therefore, we decided to test if it is possible to swap the charge distribution in β3 and β6 by mimicking the charge pattern of the gp41-1 intein onto the *Npu*DnaE intein. We introduced three lysines in Int_N_ and three glutamates in the Int_C_ of *Npu*DnaE intein (Fig. 4A). This charge swapped *Npu*DnaE intein (CS-*Npu*DnaE) could efficiently splice in *cis*, confirming that swapping these charges does not influence protein splicing in *cis* (Fig. 4B). This result is in line with the previous report in which the charge-swapping of *Npu*DnaE intein was successfully introduced into the entire *Npu*DnaE_N_ and *Npu*DnaE_C_ fragments to suppress the cross-reactivity^23^.

**Fig. 4.**
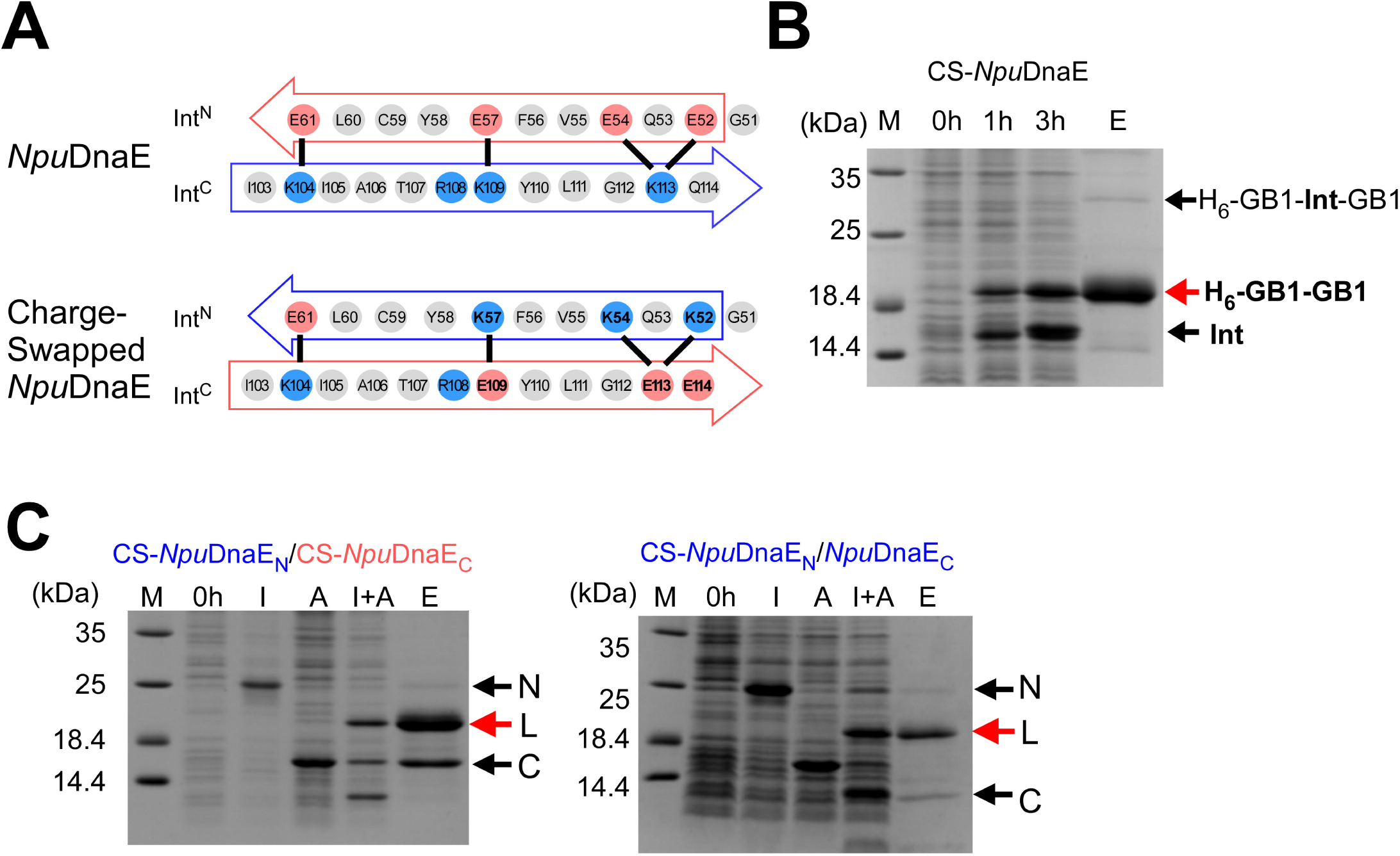
Charge engineering of the *Npu*DnaE intein. (**A**) The charge distributions of *Npu*DnaE intein and the charge-swapped *Npu*DnaE (CS-*Npu*DnaE) intein corresponding to β3 and β6 strands in the gp41-1 intein structure. (**B**) SDS-PAGE analysis of *cis*-splicing of CS-*Npu*DnaE intein. M, 0h, 3h, and E stand for molecular markers, 0 hours, 3 hours after induction, and elution from Ni-NTA spin columns. A red arrow indicates the band corresponding to the *cis*-spliced H_6_-GB1-GB1 product. (**C**) Cross-activity between split *Npu*DnaE and CS-*Npu*DnaE inteins. The left panel shows SDS-PAGE analysis of *trans*-splicing using the split CS-*Npu*DnaE intein (CS-*Npu*DnaE_N_/CS-*Npu*DnaE_C_). The right panel presents *trans*-splicing between the N-terminal fragment of the split CS-*Npu*DnaE intein (CS-*Npu*DnaE_N_) and the original C-terminal fragment of the split *Npu*DnaE intein (*Npu*DnaE_C_). N, L, and C denote the N-terminal fragment with Int_N_, the ligated product, and the C-terminal fragment with Int_C_, respectively. I, A, I+A, and E stands for IPTG induction, arabinose induction, both IPTG and arabinose induction, and elution from Ni-NTA spin columns. IPTG induction produces the N-terminal precursor fragment (N). Arabinose indication induces the protein expression of the C-terminal precursor (C). Only dual induction by IPTG and arabinose (I+A) is expected to produce the ligated product (L) by protein *trans*-splicing. The full-length gels for **B** and **C** are presented in Supplementary Figure 1.

### Orthogonality of charge swapped split inteins

The naturally split gp41-1 intein seems to be sensitive to any changes in the splice junctions as well as in its loops, which could constrain its practical applications by PTS (Fig. 2E). In contrast, the naturally split *Npu*DnaE intein and its homologs are more tolerant of sequence changes at the splice junctions, making it more suitable for protein engineering applications than the gp41-1 intein^14,32^. However, naturally split DnaE inteins from cyanobacteria are cross-reactive to each other ^18,32^. This cross-activity among naturally split DnaE inteins could limit their applications for, e.g., one-pot three-fragment ligation by PTS employing two split inteins. For such multi-fragment applications, two non-cross active (orthogonal) split inteins are required in order to suppress undesired cross-activity. Several approaches have been used to circumvent the cross-reactivity, such as utilizing different split sites of the *Npu*DnaE intein or kinetic control of two split inteins^21,22,23^. The three-dimensional structure of the gp41-1 intein revealed charge distributions different from the *Npu*DnaE intein in the corresponding β3 and β6 strands. Next, we asked if the charge network found in naturally split inteins can be responsible for the orthogonality of split inteins. We created a split intein from the charge-swapped *Npu*DnaE intein (CS-*Npu*DnaE) and tested the cross-activity of the N-terminal split intein (CS-*Npu*DnaE_N_). CS-*Npu*DnaE_N_ could still sufficiently splice with the wild-type Int_C_ of *Npu*DnaE intein (*Npu*DnaE_C_), suggesting that the charge network in the region of β3 and β6 alone cannot account for the cross-activity among the naturally split DnaE inteins (Fig.4C). Nevertheless, the charges in this region may play an important role, e.g., for making split intein fragments more soluble. This observation is consistent with the previous report that the C-terminal 16-residue fragment of *Npu*DnaE intein is sufficient for efficient *trans*-splicing of the *Npu*DnaE intein^33,34,35^.

### Engineering of orthogonal split inteins

In contrast to naturally split inteins, *cis*-splicing inteins generally possess a less pronounced charge network within the regions corresponding to β3 and β6 in the gp41-1 structure^29^. Artificially split inteins derived from *cis*-splicing inteins are often poorly soluble and might not be suitable for protein ligation because they would require unfolding/refolding processes to initiate protein *trans*-splicing^38,39^. Hence, it would be of particular interest if one could introduce the charge network similar to the one observed in naturally split inteins into the *cis*-splicing *Npu*DnaB mini-intein, which is the closest structural homolog of the gp41-1 intein and superior in tolerating sequence alterations at the splice junctions with high splicing activity^15,36,37^. We introduced five lysine residues (three in β3 and two in β6) (Fig. 5A). The charge-introduced *Npu*DnaB mini-intein (CI-*Npu*DnaB intein) was still able to splice in *cis* efficiently (Fig. 5B). As the introduced charged residues did not impair the *cis*-splicing, we derived a split intein pair from the CI-*Npu*DnaB mini-intein (CI-*Npu*DnaB_N_/CI-*Npu*DnaB_N_) by splitting at the conserved insertion site of the homing endonuclease domain^15,25^. *Cis*-splicing, *trans*-splicing of the split intein from the CI-*Npu*DnaB mini-intein became less efficient than that of the split intein derived *Npu*DnaB mini-intein (Fig. 5C and 5D). This observation suggests that the charge interactions in the β3 and β6 could play a critical role in the association of the two split intein fragments derived from *Npu*DnaB mini-intein. To confirm this hypothesis, we introduced unfavorable interactions by mutating Lys58 to Glu in β3 and Glu116 to Lys in β6. This orthogonal design of *Npu*DnaB mini-intein (Oth-*Npu*DnaB) was still able to efficiently splice in *cis* as no precursor had been left due to spontaneous splicing when expressed in *E. coli* (Fig. 5B). The efficient *cis*-splicing of Oth-*Npu*DnaB intein verifies that the introduced mutations were not detrimental to protein splicing reaction, which is an important prerequisite for the design of split inteins. We thus split Oth-*Npu*DnaB intein into a pair of two fragments of Oth-*Npu*DnaB_N_/Oth-*Npu*DnaB_C_ at the canonical split site and tested the *trans*-splicing activity (Fig. 5D). Unlike the covalently connected *cis*-splicing Oth-*Npu*DnaB intein, *trans*-splicing between Oth-*Npu*DnaB_N_/Oth-*Npu*DnaB_C_ derived from Oth-*Npu*DnaB intein was drastically impaired (Fig. 5D). Whereas the combination of CI-*Npu*DnaB_N_/Oth-*Npu*DnaB_C_ did not give any *trans*-spliced product, the pair of CI-*Npu*DnaB_N_/*Npu*DnaB_C_ could still produce the ligated product (Fig. 5E). This observation confirmed that *Npu*DnaB_C_ and Oth-*Npu*DnaB_C_ fragments have become orthogonal with CI-*Npu*DnaB_N_. Engineering of the charge network in the corresponding to β3 and β6 in the gp41-1 intein was indeed sufficient to create orthogonal split inteins from a *cis*-splicing intein, at least with the example case of the *Npu*DnaB mini-intein.

**Fig. 5.**
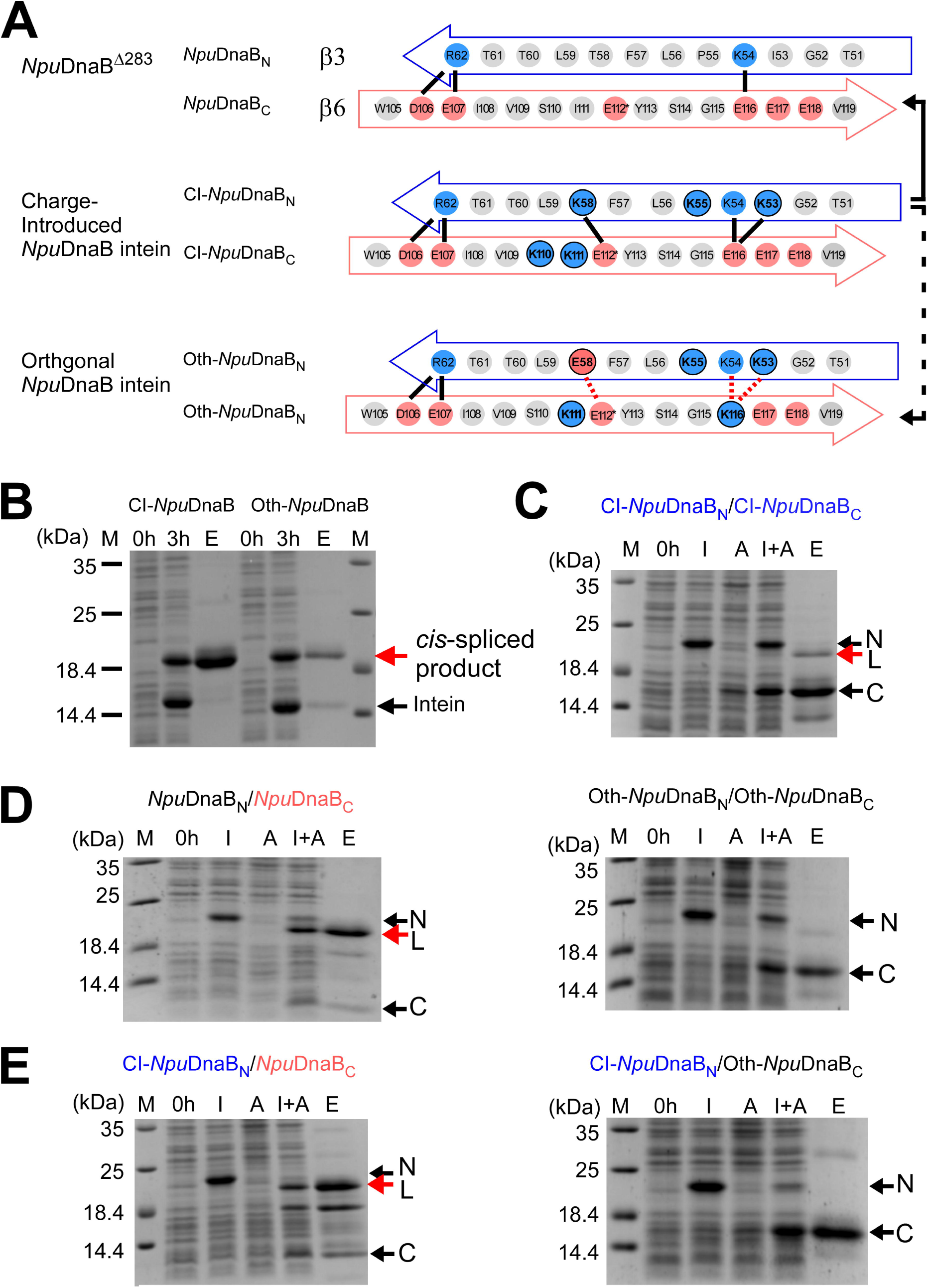
Engineering of *cis*-splicing *Npu*DnaB mini-intein toward an orthogonal pair of split inteins. (**A**) Schematic comparison between the original and engineered *Npu*DnaB mini-inteins in the regions corresponding to β3 and β6 in the gp41-1 intein structure. Solid lines indicate possible favored charge interactions. Dotted lines indicate possible unfavorable charge interactions. An arrow with a solid line indicates a charge-complemental pair for *trans*-splicing. An arrow with a broken line indicates an orthogonal pair containing unfavored charge interactions. (**B**) SDS-PAGE analysis of *cis*-splicing of the CI-*Npu*DnaB and the designed orthogonal *Npu*DnaB mini-intein. *Cis*-spliced product and excised intein are indicated by red and black arrows, respectively. M, 0h, 3h, and E above lanes indicate molecular markers, before induction, 3 hours after induction, and elution fraction from Ni-NTA columns. (**C**) The SDS-PAGE analysis of *trans*-splicing of the split version of CI-*Npu*DnaB intein, in which a split site was introduced at the canonical natural split site. (**D**) *Trans*-splicing of the split inteins derived from *Npu*DnaB intein (left) and the CI-*Npu*DnaB intein (right). (**E**) Test for orthogonality. SDS-PAGE gels show *trans*-splicing between the N-terminal fragment of CI-*Npu*DnaB mini-intein (CI-*Npu*DnaB_N_) and the C-terminal fragment of *Npu*DnaB mini-intein (*Npu*DnaB_C_) (left) and *trans*-splicing between the N-terminal fragment of CI-*Npu*DnaB mini-intein (CI-*Npu*DnaB_N_) and the C-terminal fragment derived from Oth-*Npu*DnaB mini-intein (Oth-*Npu*DnaB_C_). For panels **C**-**E**, arrows with N and C indicate the bands for the N-terminal and C-terminal precursors, respectively. L indicates the ligated product by PTS. M, 0h, I, A, I+A, and E stands for molecular markers, before induction, IPTG induction, arabinose induction, both IPTG and arabinose induction, and elution fraction from Ni-NTA spin columns, respectively. Full-length gels are presented in Supplementary Figure 2.

## Discussion

Intein-based technology gave rise to a broad range of widely applied methods in both biotechnology and basic research. Many of these methods utilize naturally occurring split inteins and their homologs because artificially split inteins derived from *cis*-splicing inteins are usually less efficient than naturally occurring split inteins and/or requiring denaturing/refolding processes due to the poorer solubility^15,24,25,38,39^. Mainly, the solubility of precursor fragments is critical for *in vitro* applications, which might obligate labor-intensive denaturing/refolding process of precursor proteins, thereby limiting their use for diverse applications. Previously, artificially splitting several *cis*-splicing inteins has not been hugely successful, resulting in less productive *trans*-splicing^15,25^. This problem has been alleviated in part by salt-inducible split inteins derived from a salt-inducible intein from extreme halophilic archaea, but yet requiring salt-induction for *trans*-splicing^40^. Having more than two robust split inteins which are not cross-active (orthogonal) could widen the applications of PTS because two orthogonal split inteins enable us to perform three-fragment ligation efficiently^20,21,22,23^. Naturally split DnaE inteins such as *Npu*DnaE intein, however, are cross-active to other naturally occurring split DnaE inteins, despite its robust splicing activity and high tolerance of variations at the splice junctions^29,32,41^. The gp41-1 intein is another fragmented split intein with robust splicing activity^19^, which could be used as an orthogonal intein with respect to other split DnaE inteins. However, the splicing activity of the gp41-1 intein turned out to be more sensitive to variations at the splice junctions (Fig. 2). We determined the crystal structure of the *cis*-splicing gp41-1 intein at 1.0 Å, which is hitherto the highest resolution available for intein structures. The structure shed light on the features common between the two naturally occurring split inteins, providing the structural basis to guide the engineering of split inteins from *cis*-splicing inteins. Both three-dimensional structures of the naturally split gp41-1 and *Npu*DnaE inteins highlighted the extended charge network on the strands corresponding to β3 and β6 in the gp41-1 intein structure, which is absent in many of *cis*-splicing inteins and could play an essential role for split inteins to be more efficient in *trans*-splicing. The charge swapping in the corresponding β3 and β6 regions in the *Npu*DnaE intein did not affect *cis*-splicing as well as *trans*-splicing, suggesting that the charge network in β3 and β6 regions alone cannot sufficiently account for the orthogonality of the naturally split *Npu*DnaE intein.

In contrast, we demonstrated that the charge engineering of split *Npu*DnaB mini-intein derived from a *cis*-splicing intein in the same β3 and β6 regions could become orthogonal. *Trans*-splicing of the *Npu*DnaB mini-intein *in vivo* can be as efficient as the *Npu*DnaE intein and has a high tolerance of variations at the splicing junction^15,36,37^. Split inteins engineered from *Npu*DnaB mini-intein are new additions to the protein engineering toolbox using protein *trans*-splicing and contribute in overcoming the junction sequence and extein dependencies currently complicating PTS applications. More than 1500 inteins or intein-like domains have been identified from the sequence databases^42^. Not all *cis*-splicing inteins could be converted into active mini-inteins by deleting the inserted homing endonuclease regions due to the mutualism developed between HINT and homing endonuclease domains^28,43^. However, a few hundred mini-inteins carrying various junction sequences in the intein database remain experimentally untested and unexplored. The common structural features found among naturally split inteins could be exploited to convert many other naturally occurring *cis*-splicing inteins into natural-like split inteins with robust *trans*-splicing activity. This process would result in the creation of new orthogonal split inteins with desired features such as optimal junction sequences and high tolerance of the foreign extein sequences, and lead to expanding the applicability of protein *trans*-splicing in protein engineering, chemical biology, and synthetic biology, particularly when scar-less protein ligation is critical for the applications.

## Methods

### Plasmid constructions

All plasmids used and designed in this study are listed and summarized in Supplemental Table S1, including the oligonucleotide sequences used. The gp41-1 intein variant for crystallization was cloned in pBHRSF38 as a SUMO fusion protein with an inactivating C1A substitution and a stop codon after the last residue of Asn125 for purification^44^. *Cis*-splicing gp41-1 intein variants with a loop or junction variations are encoded in plasmids pADHDuet21, pBHDuet37, pBHDuet321, pHBDuet021, pHBDuet087, and pHBDuet088. pHBDuet093 is a *cis*-splicing vector with the charge-swapped *Npu*DnaE intein (Supplemental Table 1). *Cis*-splicing vectors containing the charge-introduced and orthogonal *Npu*DnaB mini-intein variants are pHBDuet139 and pHBDuet140, respectively. A dual vector system using a pair of pHBDuet095 and pHBBAD106, derived from pHBDuet093, was used for testing *trans*-splicing of CS-*Npu*DnaE intein, in which the N-and C-terminal fragments can be induced with isopropyl β-D-1-thiogalactopyranoside (IPTG) and arabinose, respectively^45^. Two previously described plasmids pSADuet259 (Addgene #121910) and pSABAD250 (Addgene #45612) encoding *Npu*DnaB^Δ283^ _ΔC39_ and *Npu*DnaB _C39_, respectively, were used as a reference for *trans*-splicing of the *Npu*DnaB mini-intein^15^. Plasmid pSKBAD2 (Addgene #15335) encoding the natural NpuDnaE_C_ intein fragment was used to test orthogonality of CS-*Npu*DnaE_N_^32^. A pair of two precursor fragments with CI-*Npu*DnaB^Δ290^ _ΔC39_ (pHBDuet148, Addgene #121911) and CI-*Npu*DnaB_C39_ (pHBBAD113, Addgene #121912) were derived from plasmid pHBDuet139 (Addgene #121913). Split intein fragments derived from Oth-*Npu*DnaB, i.e., Oth-*Npu*DnaB^Δ290^ _ΔC39_ and Oth-*Npu*DnaB _C39_, were encoded in pHBDuet116 (Addgene #121915) and pHBBAD168 (Addgene #121916), respectively, which were derived from pHBDuet140 (Addgene #121914). The plasmids with Addgene numbers are available from www.addgene.org

### Protein production and purification

All recombinant proteins were produced in *E. coli* T7 Express (New England Biolabs, Ipswich, USA). For small-scale expression and purification of amounts sufficient to analyze protein splicing in *cis* and *trans*, 5 mL LB medium cultures supplemented with 25 µg mL^-1^ kanamycin, 100 µg mL^-1^ ampicillin, or both were grown at 37°C until an OD_600_ of 0.6 was reached. Cultures to express a precursor protein containing a *cis*-splicing intein were then induced with a final concentration of 1 mM IPTG for 3 hours. For co-expression of N-and C-terminal precursors for *trans*-splicing, 0.04%-arabinose induction was followed by IPTG addition with a delay of 30 min at 30°C. The co-expression lasted for a total time of 4 hours. The cell cultures were harvested by centrifugation at 4700 *xg* for 10 min, 4°C and lysed using 400 µL B-PER bacterial protein extraction reagent (Thermo Scientific, MA, USA) according to the instructions of the manufacturer. Elution fractions from IMAC purification using Ni^2+^-NTA spin columns (QIAGEN, Netherland) were analyzed by 16.5% polyacrylamide SDS-PAGE gels stained with Coomassie Blue.

Inactive gp41-1 intein with C1A mutation utilized in structural studies was produced in 2-liter LB medium supplemented with 25 µg mL^-1^ kanamycin by induction with a final concentration of 1 mM IPTG at an OD_600_ of 0.6 for 3 hours. The cells were harvested by centrifugation and lysed in Buffer A (50 mM sodium phosphate pH 8.0, 300 mM NaCl) by continuous passaging through an EmulsiFlex-C3 homogenizer (AVESTIN) at 15000 psi for 10 min, 4°C. The cell lysate was cleared by centrifugation at 38000 *xg* for 60 min, 4°C and loaded on a HisTrap column (GE Healthcare, Chicago, Illinois, United States) for purification. The protein was purified by following the two-step protocol as previously described including the removal of the hexahistidine tag and SUMO fusion domain^44^. The purified protein contained an additional sequence “SGG” as the N-terminal extein sequence of the gp41-1 intein. For crystallization, the protein was dialyzed against deionized water and concentrated to a final concentration of 45 mg/mL using an ultrafiltration device.

### Crystallization, data collection, and structure solution

Diffracting crystals of the fusion protein comprising the N-and C-terminal gp41-1 intein fragments were obtained at room temperature by mixing 100 nL concentrated protein with 100 nL mother liquid (100 mM citric acid, pH 3.5, 100 mM magnesium sulfate, and 30% w/v PEG 3350). 30% PEG 3350 was sufficient as a cryoprotectant. Data were collected at beamline i02 at Diamond Light Source (Didcot, UK) equipped with a Pilatus 6MF detector. Data were processed using HKL3000^46^ at the nominal resolution of 1.02 Å (Table 1). The structure was solved by molecular replacement with Phaser^47^ using the *Npu*DnaE intein (PDB ID: 4kl5)^48^ as the starting model. The structure was rebuilt with Coot^49^ and refined with REFMAC5^50^. Although data completeness in the outermost shell (1.04 −1.02 Å) was only 37%, the <*I/*σ*I*> ratio was quite significant at 2.8. Since the completeness in the 1.08-1.06 Å shell was 74% and <*I/*σ*I*> 3.6, we could safely claim the effective resolution of at least 1.06 Å. However, all data were used in refinement, with almost 2800 reflections present beyond this effective resolution limit.

The protein chain could be traced in the electron density without breaks for all 128 residues (125 intein residues and three residues of the amino acid sequence “SGG” preceding the first intein residue). Alternate conformations were modeled for 22 protein residues, extending to the main chain for 18 of them. A non-canonical *cis* peptide bond was modeled between Lys87 and Glu88 based on unambiguous electron density for this part of the main chain (the electron density is also unambiguous for the side chain of Lys87, whereas the side chain of Glu88 appears to be partially disordered). Final validation was performed with MolProbity^51^, showing an acceptable quality of the model (score 1.8, 35^th^ percentile). The coordinates and structure factors were deposited in the Protein Data Bank with the accession code 6qaz.

## Acknowledgments

We thank B. Haas, S. Jääskeläinen, AD. Hietikko for their technical help in the preparation of proteins and plasmids. We thank Dr. K. Kogan for his assistance at the crystallization facility. This work was supported in part by the Academy of Finland (137995, 277335), Novo Nordisk Foundation (NNF17OC0025402 to H.M.B., NNF17OC0027550 to H.I.), Sigrid Juselius Foundation and by the Intramural Research Program of the NIH, National Cancer Institute, Center for Cancer Research, as well as with Federal funds from the National Cancer Institute, NIH, under Contract No. HHSN261200800001E (to M.L.). The crystallization and NMR facilities at the Institute of Biotechnology have been supported by Biocenter Finland and HiLIFE-INFRA. The content of this publication is solely the responsibility of the authors and does not necessarily represent the official views or policies of the U. S. Department of Health and Human Services, nor does the mention of trade names, commercial products, or organizations imply endorsement by the U. S. Government.

## Author Contributions

HI designed and supervised the project; HMB, KMM, and HI performed the experiments and analyzed data; ML and AW participated in the crystallographic studies. All authors contributed to writing the manuscript.

