## Supplemental_Information for "The crystal structure of the naturally split gp41-1 intein guides the engineering of orthogonal split inteins from a *cis*-splicing intein"

#### Supplemental Table S1

Plasmids and oligonucleotides used in this study. Circularization of linear PCR products was done using Gibson-Cloning (Gibson *et al.*, 2009).

| Plasmid | Description | Reference |
|---|---|---|
| pADHDuet21 | **$P_{T7}$::$H_6$-GB1-GSY-gp41-1 intein-SSG-GB1**<br>The gp41-1 intein coding sequence was PCR-amplified from pBHDuet37 using the oligonucleotides, J557:5'-GAAGGATCCTACTGCCTGGATCTGAAAACGCAG and J558:5'-CTG-GGTACCGCTGCTGTTGTGGGTCAGAATGTC, thereby adding sequence encoding the wild-type junction sequence at the positions -1 (Y), +1 (S), and +2 (S) to the intein. The PCR product was ligated into pSKDuet16 using *Bam*HI/*Kpn*I restriction sites. The three residues of N- and C-terminal junction sequences are "GSY" and "SSG", respectively. | This work |
| pALBDuet28 | **$P_{T7}$::$H_6$-GB1-*Npu*DnaB$^{\Delta 290}$ intein-GB1**<br>An IPTG-inducible bacterial expression vector encoding the *Nostoc punctiforme* DnaB$^{\Delta 290}$ mini-intein having a non-essential endonuclease domain deleted, flanked by two GB1 domains with an N-terminal hexahistidine tag. | Aranko *et al.* (2014) |
| pBHDuet37 | **$P_{T7}$::$H_6$-GB1-EGS-gp41-1 intein-SGT-GB1**<br>The sequence encoding a fusion of the N- and C- terminal split fragments of the gp41-1 intein was codon-optimized for protein expression in *E. coli* and synthesized (IDT) with flanking *Bam*HI and *Kpn*I restriction sites and ligated into pSKDuet16 resulting in a bacterial expression vector encoding the gp41-1 intein flanked by two GB1 domains with N-terminal hexahistidine tag. The three residues of N- and C-terminal junction sequences are "EGS" and "SGT", respectively. | This work |
| pBHDuet321 | **$P_{T7}$::$H_6$-GB1-EGS-gp41-1$^{\Delta 2aa}$ intein-SGT-GB1**<br>pBHDuet321 was constructed from pBHDuet37 by inverse PCR using the oligonucleotides J269:5'-CCTGTATATTGAAGAAGGTAAAAAGATTCTGAAAATTG and J270:5'-AATCTTTT-TACCTTCTTCAATATACAGGCACATGCCC, resulting in the loop sequence "IEEG" from the residues 86-91 "VKEMML". | This work |
| pBHRSF38 | **$P_{T7}$::$H_6$-SUMO-gp41-1(C1A)**<br>The gp41-1 intein coding sequence was amplified from pBHDuet37 using the oligonucleotides I521:5'-TTGGATCCGGTGGTGCCCTGGATCTGAAAACGCAG and I522:5'-TTGGATCCGG-TGGTGCCCTGGATCTGAAAACGCAG and ligated into pHYRSF53 using *Bam*HI/*Hin*dIII, resulting in a bacterial expression vector for inactive gp41-1(C1A) | This work |

| Plasmid | Description | Reference |
|---|---|---|
| | intein with N-terminal SUMO fusion and N-terminal hexahistidine tag. | |
| pHBBAD106 | **$P_{ara}$::CS-*Npu*DnaE$_C$ intein-GB1-$H_6$**<br>The 35-residue C-terminal split fragment of the *Npu*DnaE intein carrying the charge-swapping (CS) mutations K109E, K113E, and Q114E was PCR-amplified from pHBDuet093 together with GB1 sequence using the oligonucleotides SK094:5'-TAACATATGATCAAAATAGCCAC-ACG and HK158:5'-AGAATTCCGTTACGGTGTAGGTTTTG. The PCR product was ligated into *Nde*I/*Eco*RI-digested pMHBAD14, thereby attaching a C-terminal hexahistidine tag. | This work |
| pHBBAD113 | **$P_{ara}$::CI-*Npu*DnaB$_{C39}$ intein-GB1-$H_6$**<br>Point mutations encoding S110K and I111K corresponding to the *Npu*DnaB$^{\Delta 290}$ sequence were introduced into the *Npu*DnaB$_{C39}$, C-terminal 39-residue split fragment of *Npu*DnaB mini-intein, encoded in plasmid pSABAD250 by inverse PCR using the oligonucleotides L162:5'-GGGATGAAATAGTTAAAAAGGAATATAGTGGTGAGGAAG and L163:5'-CACCACTATAT-TCCTTTTTAACTATTTCATCCCAATAAAT. | This work<br>Addgene # 121912 |
| pHBBAD168 | **$P_{ara}$::Oth-*Npu*DnaB$_{C39}$ intein-GB1-$H_6$**<br>The plasmid was created from pSABAD250 by inverse PCR using the two oligonucleotides L286:5'-GGGATGAAATAGTTTCAAAGGAATATAGTGGTAAGGAAGAAGTGTT and L287: 5'-AACACTTCTTCCTTACCACTATATTCCTTTGAAACTATTTCATCCC, bearing the I111K and E116K substitutions. | This work<br>Addgene # 121916 |
| pHBDuet021 | **$P_{T7}$::$H_6$-GB1-SGY-gp41-1 intein-SSS-GB1**<br>The gp41-1 intein coding sequence was PCR-amplified from pBHDuet37 using the oligonucleotides HB013:5'-CAAAACCTACACCGTAACGGAAGGATCCGGCTATTGCCTGGATC-TGAAAACGCAGGTG and HB014:5'-CGTTCAGGATAAGTTTGTACTGGGTACCGCTCGA-GCTGTTGTGGGTCAGAATGTCGTTC, containing the N- and C-terminal three natural extein sequences of "SGY" and "SSS". The PCR product was ligated into pBHDuet37 using *Bam*HI/*Kpn*I sites. | This work |
| pHBDuet087 | **$P_{T7}$::$H_6$-GB1-SGY-gp41-1$^{\Delta KEMM}$ intein-SSS-GB1**<br>pHBDuet087 was created by inverse PCR from pHBDuet021 using the two oligonucleotides L144:5'-TGCCTGTATGTGGGTGGTCTGAAAAAGATTCTGAAAAT and L145:5'-CTTTTTCA-GACCACCCACATACAGGCACATGCCCTC, containing the "GG" loop sequence at the natural split site to replace "KEMM" (residues 87-90). | This work |
| pHBDuet088 | **$P_{T7}$::$H_6$-GB1-EGS-gp41-1$^{\Delta KEMM}$ intein-SGT-GB1**<br>The plasmid was derived from pBHDuet37 using the two oligonucleotides L144 and L145 as | This work |

| Plasmid | Description | Reference |
|---|---|---|
| | described for pHBDuet087. | |
| pHBDuet093 | **$P_{T7}$::$H_6$-GB1-CS-*Npu*DnaE intein-GB1**<br>pHBDuet093 was constructed from pSKDuet16 by two-step inverse PCR using the two oligonucleotides L137:5'-CACGATCGCGGAAAACAAAAGGTGTTTAAGTATTGTTTGGAA and L138:5'-CCAAACAATACTTAAACACCTTTTGTTTTCCGCGATCGTGCC, containing the charge-swapping (CS) mutations E52K, E54K, and E57K. The second inverse PCR was done using the two oligonucleotides L140:5'-AATGTCATAGTCATTTTCTTCGCCTAAATATTCAC-GTGTGGCTAT and L159:5'-TAGCCACACGTGAATATTTAGGCGAAGAAAATGTCTATGA-CATTG, bearing the K109E, K113E, and Q114E mutations. | This work |
| pHBDuet095 | **$P_{T7}$::$H_6$-GB1-CS-*Npu*$DnaE_N$ intein-GB1**<br>A PCR fragment containing $H_6$-GB1-CS-*Npu*$DnaE_N$ with charge-swapping (CS) substitutions including the N-terminal His-tagged GB1 was amplified from pHBDuet093 using the oligonucleotides HB007:5'-TAATACGACTCACTATAGGGGAATTGTG and L135:5'-GTGCG-GCCGCAAGCTTAATTCGGCAAATTATCAACCCGC. The backbone vector was amplified from pHYRSF53 using J502:5'-TAAGCTTGCGGCCGCACTC and J549:5'-CCATGGTATAT-CTCCTTATTAAAG, and assembled into plasmid pHBDuet095 using Gibson-Cloning. | This work |
| pHBDuet112 | **$P_{T7}$::$H_6$-GB1-*Npu*$DnaB^{\Delta 290,I53K,P55K,T58E,S110K,I111K}$ intein-GB1**<br>Charge-introducing substitutions encoding S110K and I111K were introduced into the *Npu*$DnaB^{\Delta 290}$ mini-intein coding sequence by inverse PCR from pALBDuet28 using the oligonucleotides L162:5'-GGGATGAAATAGTTAAAAAGGAATATAGTGGTGAGGAAG and L163:5'-CACCACTATATTCCTTTTTAACTATTTCATCCCAATAAAT. Substitutions encoding I53K, P55K, and T58E were then introduced by inverse PCR using L160:5'-CGACTGGTAAAAAGAAGCTGTTTGAATTGACAACTCGATTGGGG and L161:5'-CGAGTT-GTCAATTCAAACAGCTTCTTTTTACCAGTCGAAAAAGCATT. | This work<br>Addgene #121912 |
| pHBDuet116 | **$P_{T7}$::$H_6$-GB1-Oth-*Npu*$DnaB^{\Delta 290}{}_{\Delta C39}$ intein**<br>Charge-introducing substitutions encoding I53K, P55K, and T58E were first introduced into the *Npu*$DnaB^{\Delta 290}$ mini-intein by inverse PCR on pALBDuet28 using the two oligonucleotides L160:5'-CGACTGGTAAAAAGAAGCTGTTTGAATTGACAACTCGATTGGGG and L161:5'-CGAGTTGTCAATTCAAACAGCTTCTTTTTACCAGTCGAAAAAGCATT. The 98 residue-comprising N-terminal split fragment was then amplified by PCR using HK151:5'- | This work<br>Addgene #121915 |

| Plasmid | Description | Reference |
|---|---|---|
| | TAGGATCC-GGTTGTTTAGCAGGCGATAGTC and HK297:5’-GTGAAGCTTAATTTCTTGGTAAACTGA-GATGTTCT and ligated into *Bam*HI/*Hin*dIII-digested pSKDuet01, thereby attaching N-terminal His-tagged GB1. | |
| pHBDuet139 | **$P_{T7}$::$H_6$-GB1-CI-*Npu*DnaB$^{\Delta290}$ intein-GB1**<br>*Npu*DnaB$^{\Delta290,I53K,P55K,T58E,S110K,I111K}$ residue E58 encoded in plasmid pHBDuet112 was mutated to K by inverse PCR using L284:5’-GACTGGTAAAAAGAAGCTGTTTAAATTGACAACTCGA-TTGG and L285:5’-TCGAGTTGTCAATTTAAACAGCTTCTTTTTACCAGTCGAAAAAGC resulting in *cis*-splicing charge-introduced (CI)-NpuDnaB$^{\Delta290}$ intein encoding the I53K, P55K, T58K, S110K, and I111K substitutions. | This work<br>Addgene #121913 |
| pHBDuet140 | **$P_{T7}$::$H_6$-GB1-Oth-*Npu*DnaB$^{\Delta290}$ intein-GB1**<br>pHBDuet140 was derived from pHBDuet112 by inverse PCR using the two oligonucleotides L286:5’-GGGATGAAATAGTTTCAAAGGAATATAGTGGTAAGGAAGAAGTGTT and L287: 5’-AACACTTCTTCCTTACCACTATATTCCTTTGAAACTATTTCATCCC, introducing E116K. The final gene for the Oth-*Npu*DnaB$^{\Delta290}$ intein encodes the I53K, P55K, T58E, I111K, and E116K substitutions of the *Npu*DnaB$^{\Delta290}$ mini-intein. | This work<br>Addgene #121914 |
| pHBDuet148 | **$P_{T7}$::$H_6$-GB1-CI-*Npu*DnaB$^{\Delta290}{}_{\Delta C39}$ intein**<br>The N-terminal split fragment CI-*Npu*DnaB$^{\Delta290}{}_{\Delta C39}$ intein was constructed by introducing a single E58K mutation into pHBDuet116 by inverse PCR using the oligonucleotides L284:5’-GACTGGTAA-AAAGAAGCTGTTTAAATTGACAACTCGATTGG and L285:5’-TCGAGTTGT-CAATTTAAACAGCTTCTTTTTACCAGTCGAAAAAGC. | This work<br>Addgene #121911 |
| pHYRSF53 | **$P_{T7}$::$H_6$-SUMO-*Npu*DnaE$_N$-CBD**<br>IPTG-inducible bacterial expression vector encoding hexahistidine-tagged fusion protein of SUMO, the native split N-terminal *Nostoc punctiforme* DnaE intein, and CBD. | Addgene #64696 |
| pMHBAD14 | **$P_{ara}$::*Npu*DnaE$_C$-GB1-$H_6$**<br>An arabinose-inducible bacterial expression vector encoding the C-terminal native split fragment (35 residues) of the *Nostoc punctiforme* DnaE intein with C-terminal GB1 and hexahistidine tag fusion. | Addgene #42304 |
| pSABAD250 | **$P_{ara}$::*Npu*DnaB$_{C39}$-GB1-$H_6$**<br>An arabinose-inducible bacterial expression vector encoding the C-terminal artificial split fragment (39 residues) of the *Nostoc punctiforme* DnaB intein with C-terminal GB1 and hexahistidine tag fusion. | Addgene #45612 |

| Plasmid | Description | Reference |
|---|---|---|
| pSADuet259 | **$P_{T7}$::$H_6$-GB1-*Npu*DnaB$^{\Delta 290}{}_{\Delta C39}$**<br>IPTG-inducible bacterial expression vector encoding the N-terminal artificial split fragment (residues 1-98) of the *Nostoc punctiforme* DnaB intein with N-terminal GB1 and hexahistidine tag fusion. | Addgene #121910 |
| pSKBAD2 | **$P_{ara}$::*Npu*DnaE$_C$-GB1**<br>Arabinose-inducible bacterial expression vector encoding the C-terminal native split fragment (35 residues) of the *Nostoc punctiforme* DnaE intein with C-terminal GB1. | Addgene #15335 |
| pSKDuet01 | **$P_{T7}$::$H_6$-GB1-*Npu*DnaE$_N$**<br>IPTG-inducible bacterial expression vector encoding the N-terminal split fragment (102 residues) of the *Nostoc punctiforme* DnaE intein with N-terminal GB1 and hexahistidine tag fusion. | Addgene #12172 |
| pSKDuet16 | **$P_{T7}$::$H_6$-GB1-*Npu*DnaE intein-GB1**<br>IPTG-inducible bacterial expression vector encoding a fusion protein of the split DnaE intein fragments of *Nostoc punctiforme* flanked by two GB1 domains with an N-terminal hexahistidine tag. | Addgene #41684 |

Abbreviations: CBD, chitin binding domain; DnaB, bacterial helicase; DnaE, catalytic α subunit of DNA polymerase III; GB1, B1 domain of the *Streptococcus sp.* IgG binding protein G; $H_6$, hexahistidine tag; IPTG, isopropyl β-D-1-thiogalactopyranoside; SUMO, yeast small ubiquitin-like modifier domain.

#### Fig. 2E

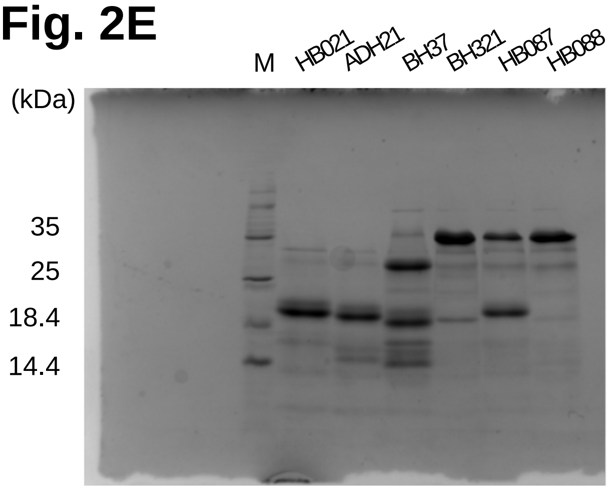


#### Fig. 4B

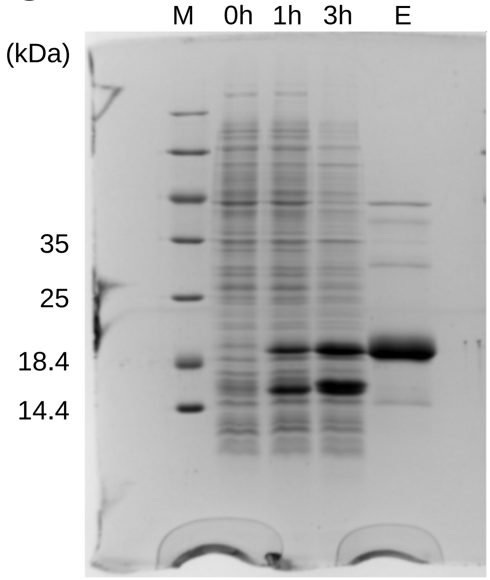


#### Fig. 4C

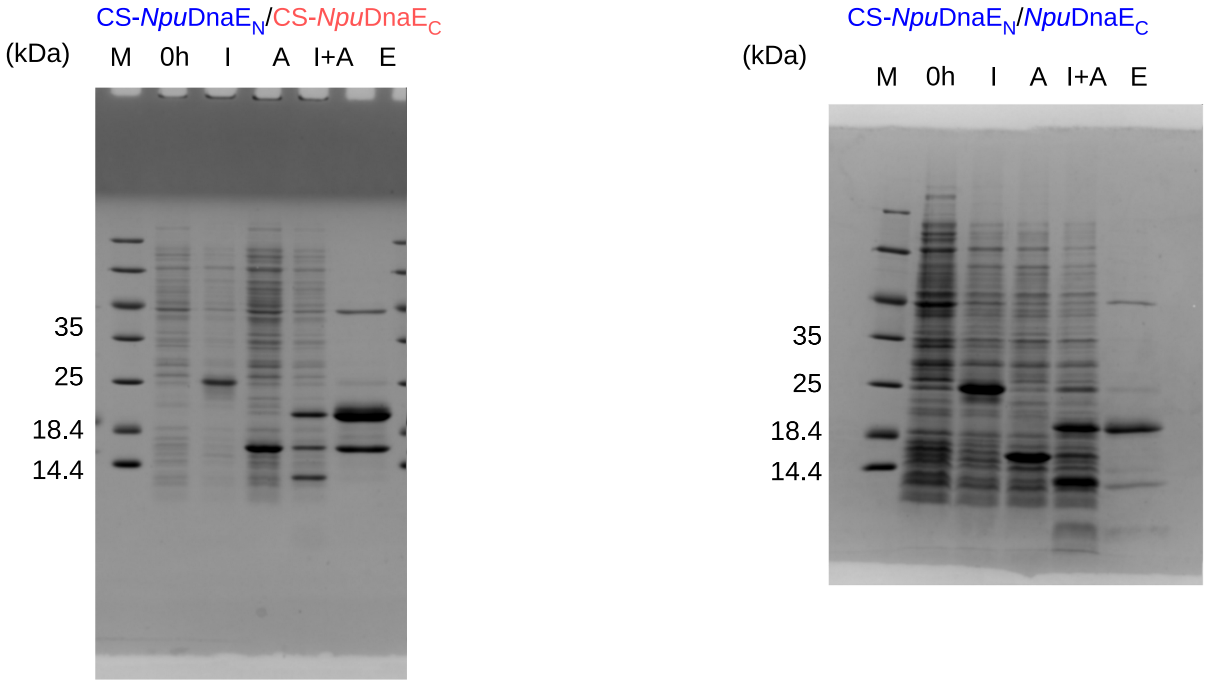


Supplemental Figure S1

### Fig. 5B

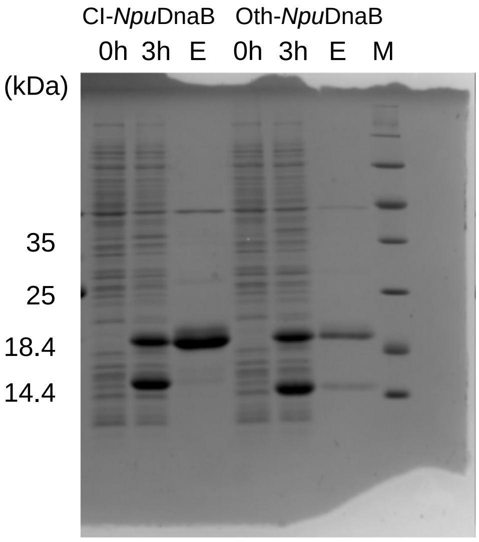


### Fig. 5C

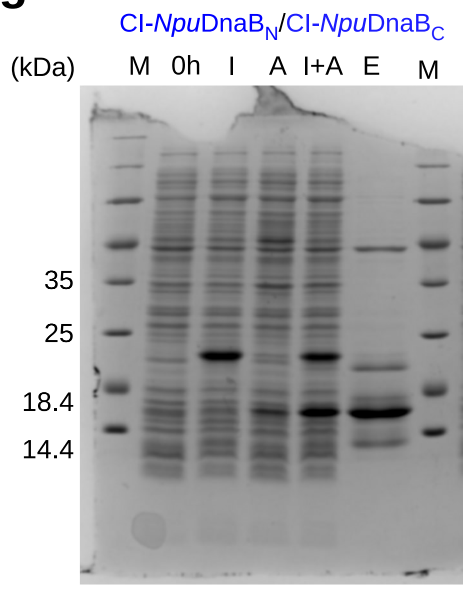


### Fig. 5D

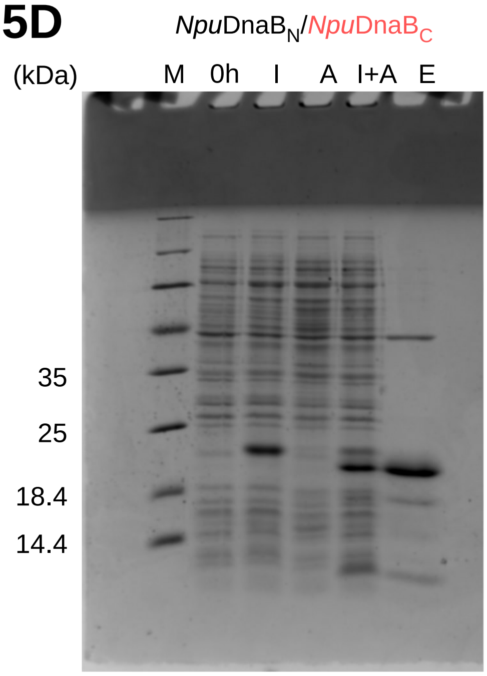


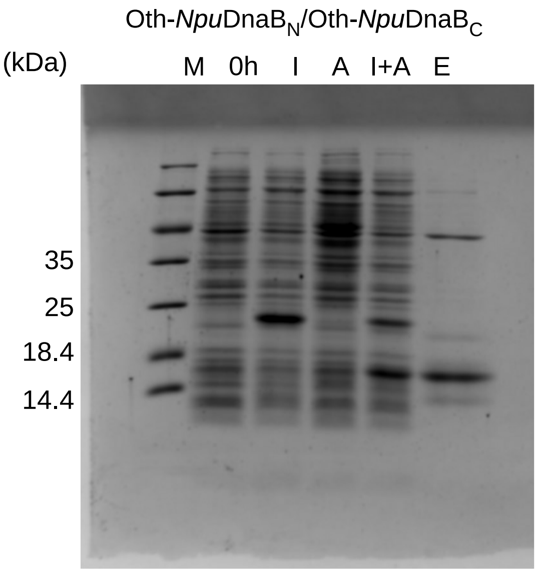


### Fig. 5E

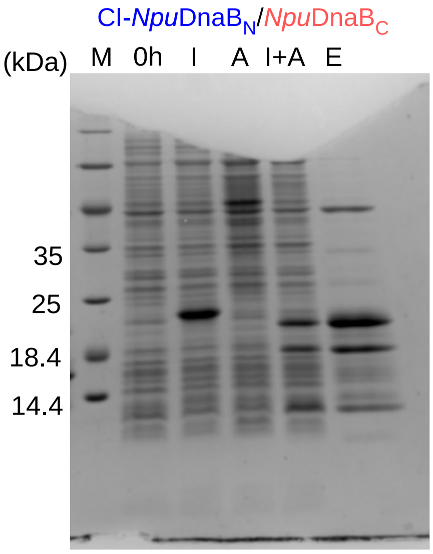


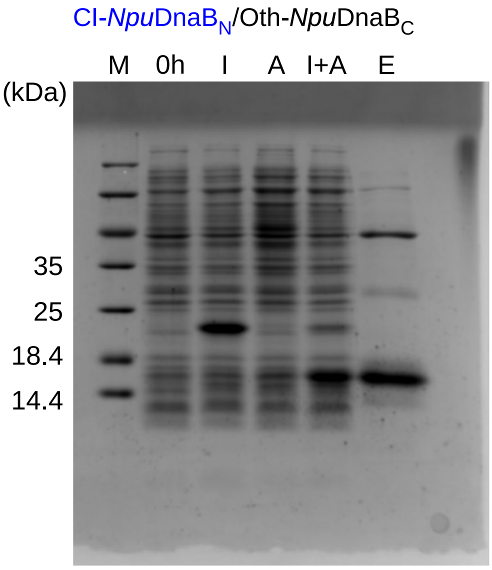


Supplemental Figure S2
